# Malaria parasite centrins assemble by Ca^2+^-inducible condensation

**DOI:** 10.1101/2022.07.26.501452

**Authors:** Yannik Voß, Severina Klaus, Nicolas P. Lichti, Markus Ganter, Julien Guizetti

## Abstract

Rapid proliferation of the malaria-causing parasite *Plasmodium falciparum* in the human blood depends on a particularly divergent and acentriolar centrosome, which incorporates several essential centrins. Centrins are small calcium-binding proteins that have a variety of roles and are universally associated with eukaryotic centrosomes. Their precise mode of action, however, remains unclear. In this study calcium-inducible liquid-liquid phase separation is revealed as an evolutionary conserved principle of assembly for *Plasmodium* and human centrins. Furthermore, the disordered N-terminus and calcium-binding motifs are defined as essential features for reversible biomolecular condensation and demonstrate that certain centrins can co-condensate. In vivo analysis using live-cell STED microscopy shows liquid-like dynamics of centrosomal centrin. Additionally, implementation of an inducible protein overexpression system reveals concentration-dependent formation of extra-centrosomal centrin assemblies with condensate-like properties. The timing of foci formation and dissolution suggests that centrin assembly is regulated. This study thereby provides a new model for centrin accumulation at eukaryotic centrosomes.

## Introduction

Malaria-causing parasites are divergent unicellular eukaryotes and still cause the death of over 600.000 people per year ^1^. To proliferate in the red blood cells of their human host, they use an unconventional cell division mode called schizogony ^2–5^. It consists of multiple asynchronous rounds of closed mitosis followed by a final round of budding, releasing 12-30 daughter cells from the bursting host cell ^6^ (Figure 1A). The high number of progeny promotes rapid proliferation and is therefore directly linked to the severity of a disease ^7^. Nuclear multiplication requires formation and duplication of the parasite microtubule organizing center (MTOC), also called centriolar plaque, which significantly differs from the highly structured spindle pole body of yeast and the centriole-containing mammalian centrosome ^8, 9^. Centriolar plaques consist of an amorphous chromatin-free intranuclear compartment from where all mitotic spindles originate, which connects through the nuclear envelope to a protein dense extranuclear compartment, where centrins localize ^10–14^. Centrins have various proposed functions and are one of the most widely conserved component of eukaryotic MTOCs ^9, 15–17^. In yeast, they localize to the half-bridge of spindle pole bodies, where they are required for duplication ^18^. In mammalian cells, centrins are found inside the centrioles and are implicated in the function of centrosomes ^19–22^. How such a small and well-conserved structural protein functions within those divergent sub-cellular contexts is still unclear. Centrins contain four EF-hand (EFh) domains that can chelate calcium with high affinity, causing conformational changes ^23, 24^. Calcium-dependent self-interaction of centrins has been documented in a range of eukaryotes including for human centrin 2 (HsCen2) ^23, 25–28^, although the nature of this interaction is unknown. While yeasts encode one centrin ^13^, humans have four centrins but only HsCen2 and 3 are directly associated with the centrosome in all tissues ^21, 29^. The centrin protein family in *Plasmodium* spp. has also expanded to four members (Figure 1B), of which three are likely essential in the blood stage of infection ^12, 13, 30, 31^. While the C-terminal sequence containing the EFh domains is homologous, their N-terminus is more variable and can contain phosphorylation sites ^32–35^. Their relocalization from the cytoplasm to the centriolar plaque at the onset of schizogony has been suggested ^11–13^. Which mechanisms drive centrin accumulation in the context of divergent eukaryotic centrosomes is unclear.

**Figure 1.**
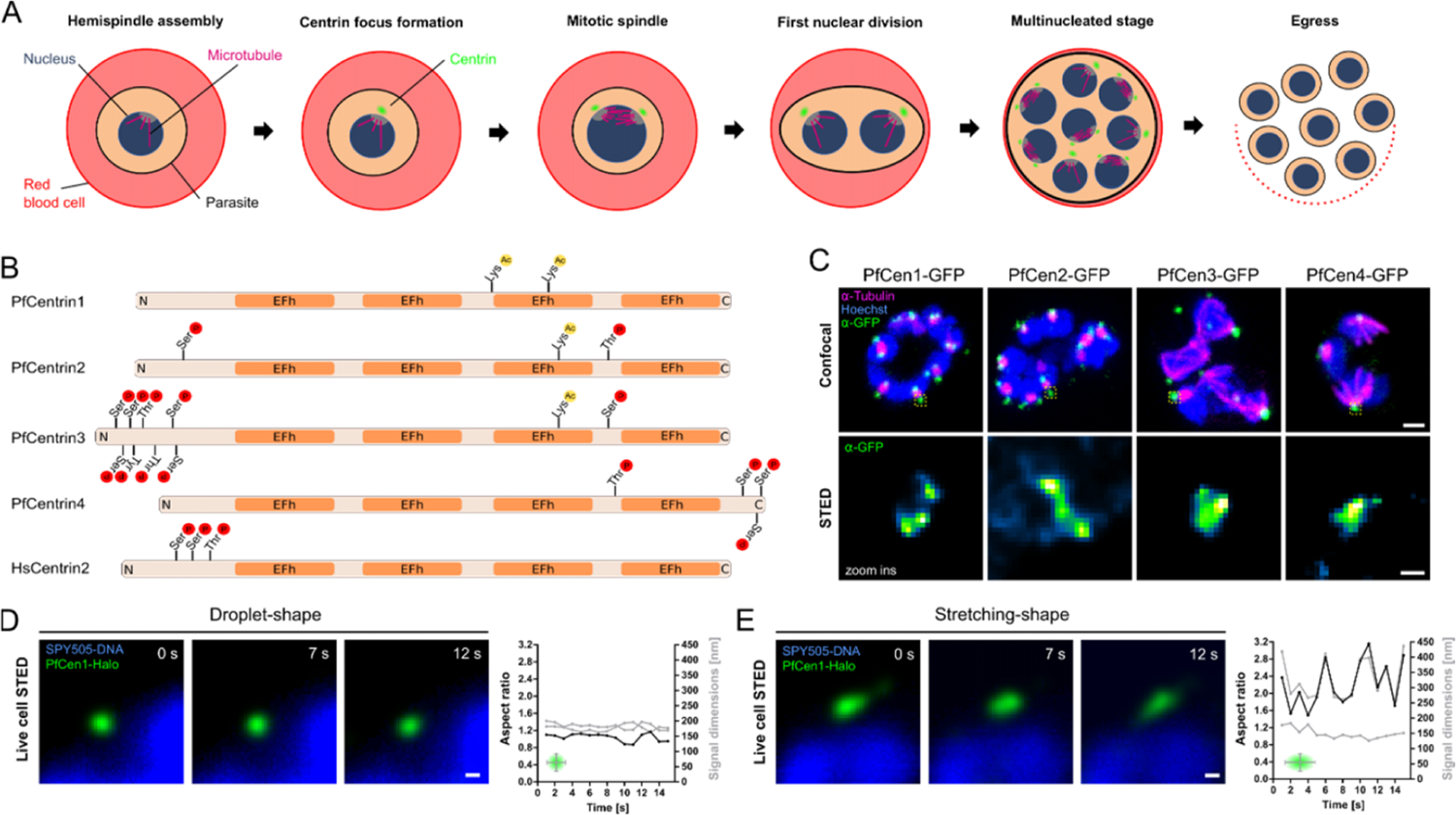
PfCen1-4 localize to centriolar plaque and can display liquid-like dynamics. (A) Schematic of nuclear and cell division during asexual blood stage schizogony (B) Schematic of PfCen1-4 and HsCen2 indicating reported post-translational modifications and EFh domains. (C) Immunofluorescence staining of tubulin and PfCen1-4-GFP in parasite strains. All images are maximum intensity projections (MIP). DNA stained with Hoechst. Scale bars; confocal, 1 µm, STED, 100 nm. (D-E) STED time lapse of centriolar plaque region of parasites expressing PfCen1-Halo labeled with MaP-SiR-Halo dye. DNA stained with SPY505-DNA. Scale bar, 100 nm. Quantification of ratio (black line) between height and width (grey lines) of the PfCen1 signal, as indicated in the small schematic.

## Results

### All *Plasmodium* centrins localize to the centriolar plaque where they can undergo dynamic rearrangement

To assess the localization of PfCen1-4, we episomally expressed GFP-tagged proteins (Supplementary Figure S1) in cultured *P. falciparum* blood stage parasites, as they did not tolerate any endogenous tagging ^12^. PfCen1-4 localized to the centriolar plaque during schizogony (Figure 1C) and Stimulated Emission Depletion (STED) microscopy showed heterogenous shapes of the fluorescent signal. To address whether this heterogeneity might be the result of a dynamic reorganization of centrin within this diffraction-limited region, we implemented live cell STED using a parasite line expressing PfCen1 tagged with Halo (Supplementary Figure S1) and labeled with the MaP-SiR-Halo fluorogenic dye ^36^ (Figure 1D-E). This either revealed ‘wobbling’ droplet-like shapes (Movie S1), or structures which seemed to stretch and retract along a defined axis (Movie S2), as the difference in aspect ratio suggest (Figure 1D-E). These dynamics were reminiscent of biomolecular condensates ^37^, concentrated protein droplets that form by liquid-liquid phase separation (LLPS), which has emerged as a biological principle explaining the formation of membrane-less organelles, including centrosomes ^38^.

### *Plasmodium* and human centrins can undergo calcium-dependent liquid-liquid phase separation

To verify whether centrins can phase-separate, we expressed PfCen1-4 and HsCen2 in *E. coli* (Supplementary Figure S2) and imaged concentrated recombinant protein solutions. After addition of calcium, we observed rapid droplet formation for PfCen1, PfCen3, and HsCen2 (Figure 2A, Movie S3-5). The droplets fused and showed surface wetting, both hallmarks of LLPS ^39^. Addition of EDTA led to instant dissolution of the droplets, except for PfCen3 where non-coalescing droplets remained (Figure 2A, Movie S6). To quantify centrin phase separation kinetics, we measured the turbidity of the solutions by light scattering at standardized concentrations (Figure 2B). Upon calcium addition, we observed a rapid increase in turbidity that was caused by protein droplet formation (Supplementary Figure S3A) in PfCen1, PfCen3, and HsCen2 solutions (Figure 2B). Addition of Mg^2+^ as an alternative bivalent cation demonstrated that the effect was calcium-specific (Supplementary Figure S3B). GFP-tagging of PfCen1 did not disrupt LLPS (Supplementary Figure S3C), which suggests that this property can be preserved in parasite lines episomally expressing tagged PfCen1 (Figure 1B-D). Since PfCen3 condensation was not completely reversible, we added EDTA at earlier timepoints and found that the irreversible fraction increased over time (Supplementary Figure S4A), suggesting a maturation of the biomolecular condensate towards a more solid or gel-like state ^40^. In contrast, maturation in PfCen1 condensates was not observed within the first 3 hours after induction (Supplementary Figure S4B). Taken together this demonstrates that centrins have differential capacity to undergo calcium-inducible LLPS.

**Figure 2.**
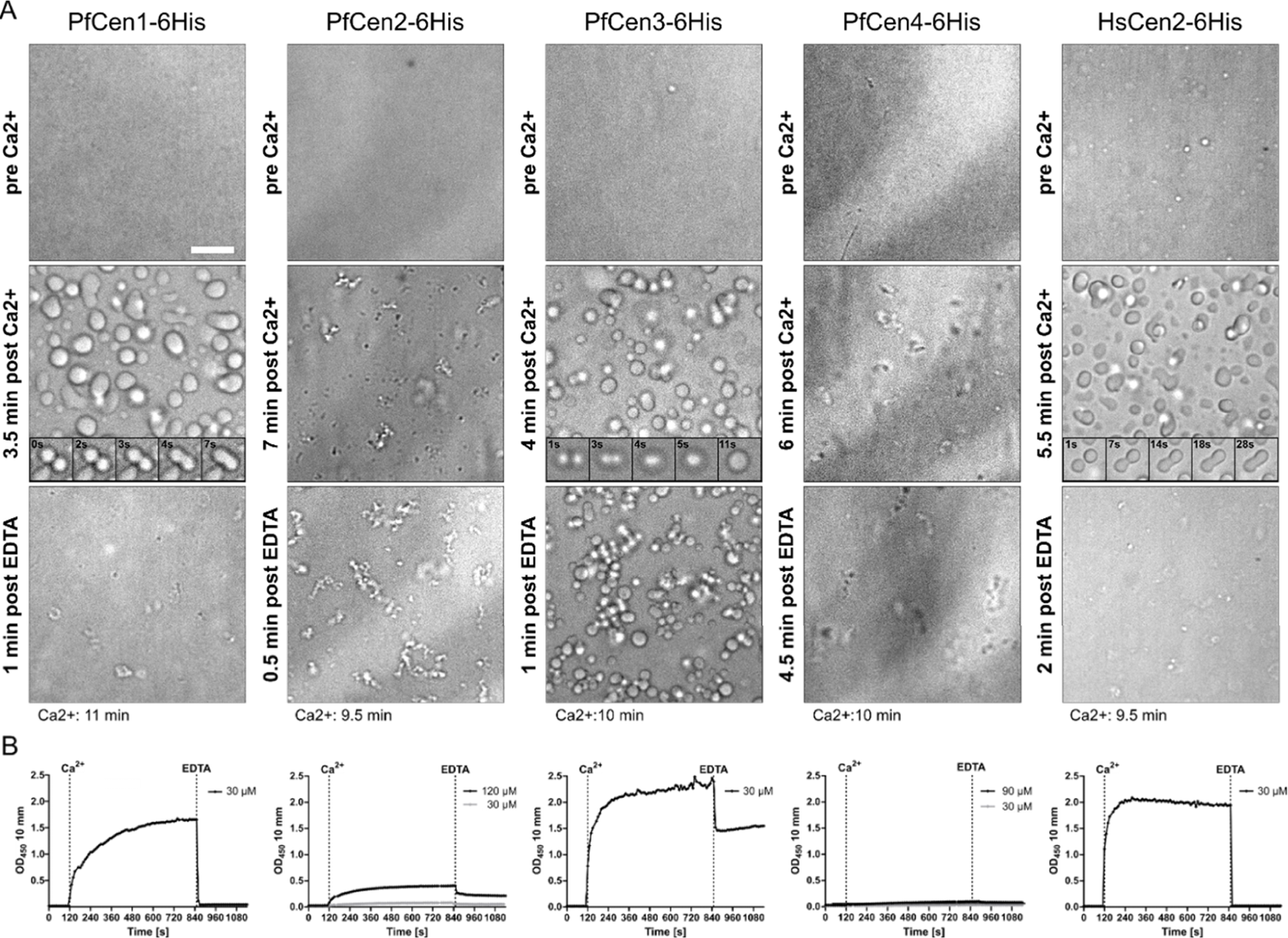
PfCen1, 3 and HsCen2 undergo calcium-dependent and reversible LLPS in vitro. (A) Widefield image of highly concentrated recombinant centrin solutions to promote high droplet density of PfCen1 (200 µM), PfCen2 (120 µM), PfCen3 (193 µM), PfCen4 (120 µM), HsCen2 (200 µM) before, after 2 mM CaCl_2_ addition, and after 10 mM EDTA addition. Scale bars, 10 µm. Inlays show time lapse images of droplet fusion events. (B) Turbidity measurements in centrin solutions at 30 µM (and higher concentration where indicated) during calcium addition followed by EDTA. Conditions: 50 mM BisTris (pH 7.1) at 37°C.

### Intrinsic disorder in the N-terminal sequence of centrins predicts self-assembly

A common feature of phase-separating proteins are intrinsically disordered regions (IDRs) ^37^, which we analyzed for the different centrins. Indeed, the prediction tool IUPred3 showed high IDR probabilities within the N-termini of PfCen1, PfCen3, and HsCen2, but not for PfCen2 and PfCen4 (Supplementary Figure S5A). Hence, presence of a N-terminal IDR predicts LLPS for centrins in the tested cases. Analysis for IDRs using IUPred3 in centrins from a range of eukaryotes suggests that most of those species encode at least one IDR-containing centrin (Supplementary Figure S5B). For some of those centrins from various distant eukaryotes, ‘polymer-like’ behavior was previously noted in vitro, further suggesting that LLPS might be an evolutionary conserved feature of a subset of centrins^25–27^.

### Phase-separating centrins can co-condensate in vitro

Interactions between centrins in malaria parasites has already been demonstrated ^12^ and is confirmed by our localization data (Figure 1B). To see whether different centrins could phase separate together, we first determined the saturation concentration, i.e., the concentration threshold at which a protein transitions from a one-phase to a two-phase regime, for PfCen1 and 3. Under the conditions presented here, we found the concentration to be around 10 µM and detected no LLPS even when increasing molecular crowding by adding 20 µM of BSA (Figure 3A-B). When, however, replacing BSA with the other centrin, we observed clear LLPS at the same total protein concentration demonstrating that PfCen1 and 3 promote each other’s condensation (Figure 3C). This effect is due to co-condensation into joint PfCen1-PfCen3 droplets, as shown by incorporation of PfCen1-GFP into preformed PfCen3 droplets (Figure 3D).

**Figure 3.**
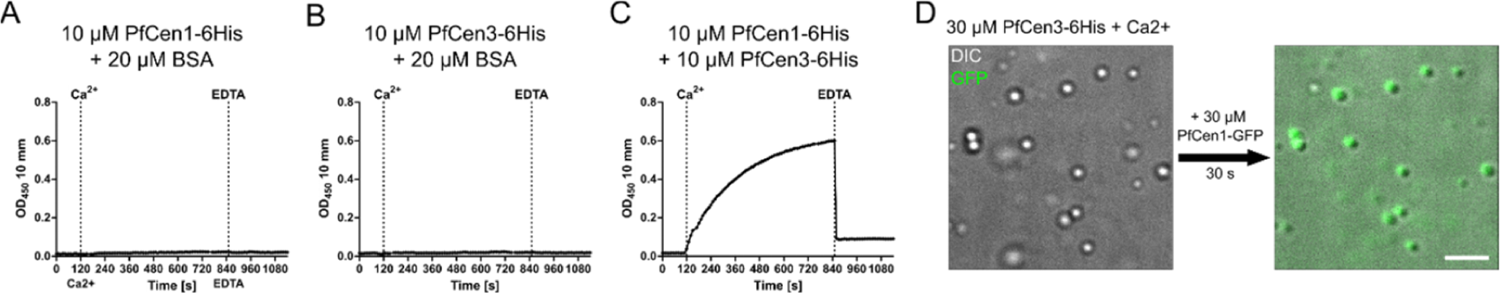
PfCen1 and PfCen3 interact through co-condensation. (A-C) Turbidity measurements of recombinant PfCen1 and PfCen3 at individually subcritical concentrations either independently or as a mixture during addition of calcium followed by EDTA. Individual centrins were supplemented with BSA to a consistent total protein concentration of 20 µM. (D) Brightfield and fluorescence imaging of preformed 30 µM PfCen3-6His protein droplets in presence of Ca^2+^ before and 30 s after addition of recombinant PfCen1-GFP-6His at 30 µM. Focus was adjusted between timepoints. Scale bar, 10 µm. All conditions: 50 mM BisTris (pH 7.1) at 37°C.

### Centrin overexpression causes premature formation of in vivo assemblies with condensate-like properties

While live cell STED hinted at liquid-like dynamics (Figure 1C-D), we wanted to investigate further whether condensation of centrins can occur in vivo. Since phase separation is concentration-dependent, we aimed to increase centrin levels to test whether we can induce additional formation of condensates in parasites. To exert control over the expression levels of PfCen1-GFP than with the classical overexpression vector (Supplementary Figs. S1, S6A), we designed a novel *P. <u>F</u>alciparum* Inducible <u>O</u>verexpression (pFIO) plasmid to be transfected in a DiCre recombinase expressing acceptor strain (Figure 3A). Upon addition of rapamycin the dimerized recombinase excises the first open reading frame, placing the gene of interest in front of the promoter. We designed a version with the medium strength *hsp70* promoter fragment (pFIO) and the stronger *hsp86* promoter (pFIO+), for which we confirmed a strong increase in median expression levels upon induction (Supplementary Figure S6B). In centrin-overexpressing cells, we frequently observed accumulations of PfCen1-GFP signal that were not associated with any spindle structure or the nucleus. Those Extra-Centrosomal Centrin Accumulations (ECCAs) occurred in 56% (n = 54) of cells carrying the medium strength promoter (Figure 4B) and in 97% (n = 60) of cells with the strong promoter (Figure 4C). The number of ECCAs correlated positively with total cell fluorescence intensity (Supplementary Figure S7). Time-lapse microscopy of overexpressed PfCen1-GFP showed a homogenous cytoplasmic distribution prior to assembly into ECCAs, sometime before the formation of the first mitotic spindle, or at MTOC foci (Figure 4D, Movie S7). Centrin accumulation by biomolecular condensation predicts that once a critical concentration threshold is crossed, additional centrin would only accrue into the condensate fraction. Machine-learning-based image analysis indeed showed that PfCen1-GFP concentration in the cytoplasm remained stable, while the fraction of PfCen1-GFP in foci increased once their formation started (Figure 4E). The partition coefficient of centrin signal between cytoplasmic and foci fraction was stable around 4.0 until it slightly increased to 5.3 after mitotic spindle formation (Supplementary Figure S8). At the end of schizogony, centrin foci disappeared, while the total centrin signal stayed constant, which indicates that centrin assemblies are dissolved (Figure 4E). Under normal conditions, centrin localizes to the centriolar plaque at the onset of schizogony as marked by the formation of the first mitotic spindle ^11^. When overexpressed, PfCen1-GFP coalescence into ECCAs and at the centriolar plaque occurred prematurely but was still temporarily linked to the entry into the schizont stage (Figure 4F). The timing of both ECCA and centriolar plaque foci formation was, however, concentration-dependent, which can best be explained reaching the critical concentration for schizogony-associated condensation earlier ^11^. Even ECCAs often formed first at the centriolar plaque before detaching, which further suggests an involvement of schizogony-specific nucleation factors (Movie S8). ECCAs and centrosomal centrin foci were negative for the protein aggregate stain Proteostat (Figure 4G) ^41^. Taken together these observations argue for a regulated condensation being part of the mechanism of centrin assembly at the centriolar plaque.

**Figure 4.**
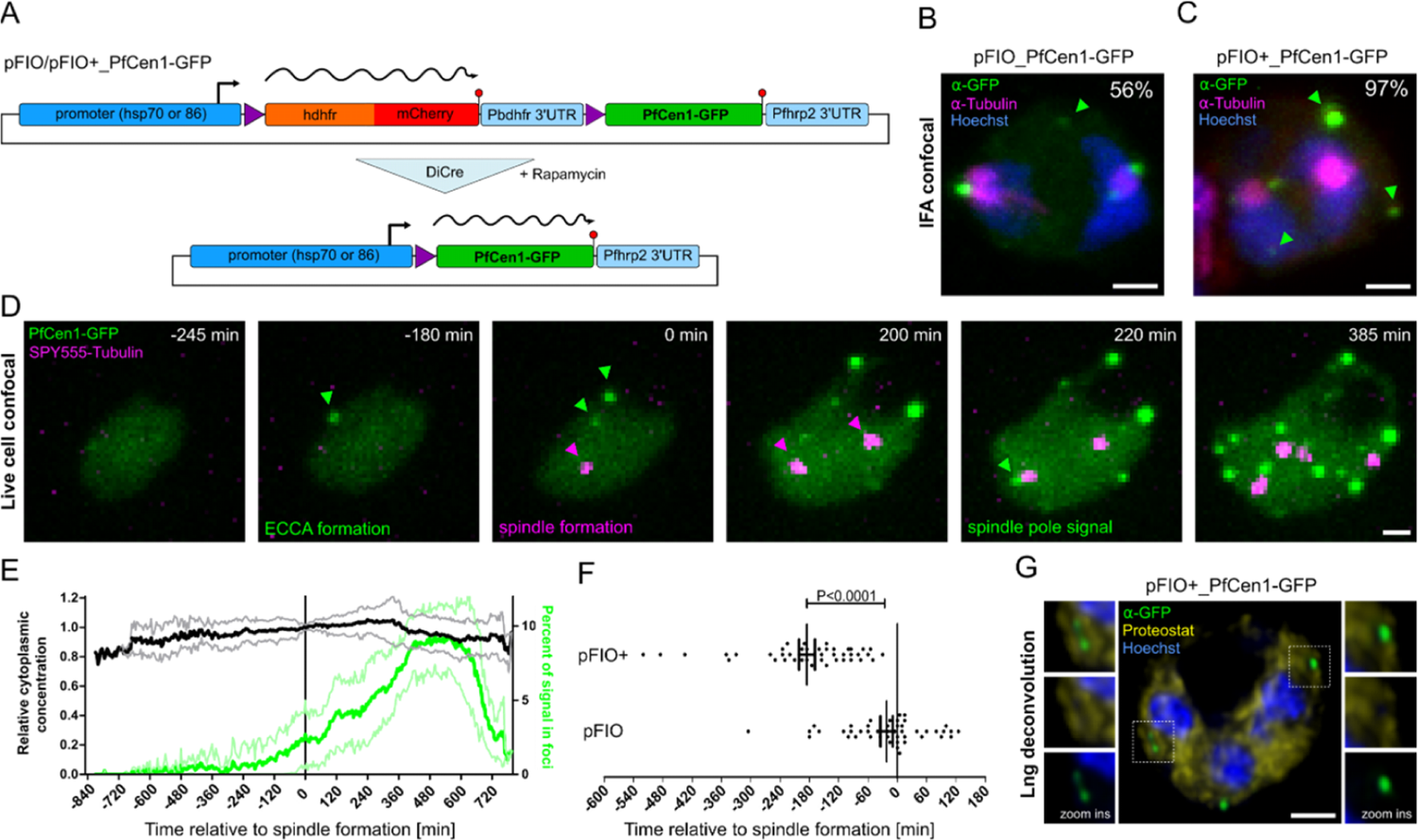
PfCen1 displays condensate-like properties in parasites. (A) Schematic of pFIO plasmids during DiCre-dependent recombination. (B) Immunofluorescence of tubulin and GFP in parasites moderately overexpressing PfCen1-GFP (pFIO). Percentage of cells containing ECCAs (arrows) indicated. (C) as in (B) but for cell strongly overexpressing PfCen1-GFP (pFIO+) (D) Time lapse images of parasites overexpressing PfCen1-GFP from pFIO+ and labeled with microtubule live stain SPY555-Tubulin (E) Normalized mean cytoplasmic PfCen1-GFP fluorescence intensity over time and share of PfCen1-GFP fluorescence signal contained within foci with standard deviation. n ≤ 33. (F) Quantification of timepoint of PfCen1-GFP foci appearance relative to first mitotic spindle formation detected by SPY650-tubulin comparing pFIO/pFIO+; standard error of the mean, two-tailed t-test, n_1_ = 46, n_2_ = 45. (G) Proteostat and PfCen1-GFP live cell staining. All images are MIP. DNA stained with Hoechst. Scale bars; 1 µm.

### PfCen1 requires its N-terminus and calcium-binding activity for condensation

To strengthen the previous conclusions, we set out to functionally interfere with LLPS in vitro and in vivo. When deleting the IDR-containing N-terminus from recombinant PfCen1, we observed strongly reduced LLPS and merely detected some aggregation (Figure 5A). LLPS was not rescued by replacement with the IDR-free N-terminus of PfCen4, neither was the N-terminus of PfCen1 sufficient to confer LLPS to a chimeric PfCen4 (Supplementary Figure S9). To test the role of EFh domains, we introduced four critical D to A mutations in the Ca^2+^-binding pocket shown to strongly inhibit Ca^2+^-binding ^42^, to generate the EFh-dead mutant, which abolished LLPS (Figure 5B). To test the significance of the IDR in vivo, we overexpressed the N-terminal deletion mutant, observing ECCAs in only 31% (n = 55) of cells (Figure 5C), as compared to 97 % upon pFIO+ overexpression of PfCen1 (Figure 4C). The EFh-dead PfCen1 mutant (Figure 5D), lacked any discernible formation of foci in vivo, suggesting a critical role for Ca^2+^ responsiveness for targeting and condensation. Together this suggests that active EFh-domains and a disordered N-terminus are essential but not sufficient for calcium-induced phase separation of centrins, with alterations inhibiting LLPS in vitro also affecting foci formation in vivo. However, deletion of the N-terminus retains some targeting activity while EFh-dead PfCen1-GFP mutants stay completely soluble.

**Figure 5.**
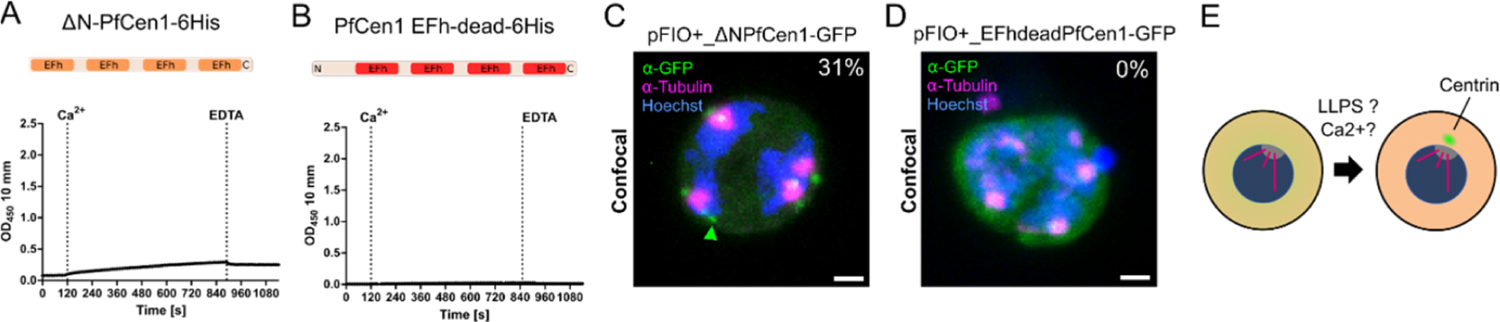
PfCen1 phase separation depends on its N-terminus and calcium binding. (A-B) Turbidity measurements in recombinant protein solutions at 30 µM of PfCen1 mutants lacking N-terminus or with disabled EFh domains, respectively, during addition of calcium followed by EDTA. Conditions: 50 mM BisTris (pH 7.1) at 37°C. (C-D) Immunofluorescence staining of tubulin and GFP in parasites overexpressing N-terminal deletion and non-Ca^2+^-binding mutant of PfCen1-GFP from pFIO+. Percentage of cells containing ECCAs (arrows) indicated. All images are MIP. DNA stained with Hoechst. Scale bars; 1 µm. (E) Our findings suggest that centrin accumulation at the centriolar plaque from a pre-mitotic diffuse cytoplasmic pool depends on LLPS, which might be promoted by cellular signaling events like calcium levels and a nucleating factor at the centrosome. Parasite is shown without host red blood cell for clarity.

## Conclusion

Centrins are conserved in all analyzed eukaryotes and implicated in a wide range of processes, which require their assembly ^23^. Our study in an early branching eukaryote revealed calcium-regulated phase separation as a new centrin assembly principle. The reported increase in intracellular calcium associated with schizogony could therefore help to promote timely centrin accumulation at the MTOC ^43^. Our data, which shows centrin assembly in vivo to be spatially and temporally linked to schizogony, however, suggests the presence of other yet to be discovered factors promoting assembly (Figure 5E). Phase condensation can further explain how the centriolar plaque remains ‘fluid’ enough to allow splitting during subsequent nuclear division cycles ^11, 14^. Biomolecular condensation and maturation have been ascribed to various centrosomal proteins ^38^, but remains controversial as the dominant principle for biogenesis of membraneless organelles ^44^. Our study suggests a combination of cell cycle-regulated nucleation with LLPS as a driver for the subsequent accumulation of centrosomal proteins.

## Materials and Methods

### Cloning of plasmids

All plasmids were assembled via Gibson assembly using the HiFi DNA Assembly mix (New England Biolabs), unless stated otherwise. Correct sequences were confirmed via Sanger sequencing (Eurofins Genomics, SupremeRun). PCRs were performed using Phusion polymerase (New England Biolabs). Details on primer sequences, templates, and synthetic oligos or genes are provided in Table S1 and S2. Plasmids were amplified in chemically competent 5-alpha F’Iq *Escherichia coli* cells (New England Biolabs) after 42°C heat shock transformation and extracted using the GenElute HP Plasmid Miniprep Kit (Sigma-Aldrich, Merck). To generate pARL-PfCen1-GFP, - PfCen2-GFP, and PfCen4_GFP from pARL-PfCen3-GFP (kindly provided by Tim Gilberger), the PfCen3 sequence was removed via restriction digest with KpnI-HF and AvrII. It was replaced with PfCen1, PfCen2, or PfCen4 sequences PCR amplified via the primer pairs P1+2, P3+4, or P5+6, respectively. To generate pARL-PfCen1-Halo the GFP sequence was removed from pARL-PfCen1-GFP via restriction digest with AvrII and PstI-HF. It was replaced with the Halo sequence PCR amplified via primer pair P7+P8. To generate pFIO pARL-Cen3-GFP was digested with NotI-HF and HindIII-HF, leaving the hrp2 3’UTR of the hDHFR (human dihydrofolate reductase) resistance cassette. Pfhsp70 5’UTR and hDHFR sequences were PCR amplified via primer pairs P9+P10 and P11+12 respectively and inserted. This created an intermediate plasmid, which was digested with BamHI-HF to insert an mCherry and GFP sequence PCR amplified via primer pairs P14+15 and P16+17 respectively. This plasmid was then digested with SalI-HF to insert a PbDHFR 3’UTR sequence PCR amplified via primer pair P18+19 to create the basic pFIO_GFP. To generate pFIO+_GFP the promotor was replaced by digestion with NotI-HF and BamHI-HF (which also removed the hDHFR sequence) and integration of Pfhsp86 5’UTR and hDHFR sequences PCR amplified using primer pairs P20+21 and P11+P13 respectively. To generate pFIO/pFIO+_Cen1-GFP, and pFIO+_ΔN-PfCen1-GFP, previously generated pFIO_GFP and pFIO+_GFP were digested with MluI-HF and XhoI to remove the GFP open reading frame (ORF), which was replaced with a PfCen1-GFP or ΔN-PfCen1-GFP ORF PCR amplified using primer pairs P22+17 or P23+17 respectively. pFIO+_ EFh-dead-Cen1-GFP, and_HsCen2-GFP were generated by removing the Cen1 ORF in pFIO+_Cen1-GFP by digestion with MluI-HF and AvrII, and replacing it with synthesized genestrings (Thermo Fisher) encoding EFh-dead-PfCen1, where aspartate 37, 73, 110, and 146 were mutated to alanine, and HsCen2 codon optimized for *P. falciparum* (oligo 1 and 2). All plasmodial vectors contained a hDHFR selection cassette conveying resistance to WR99210. pET-28 constructs were generated by digesting pET-28a(+) with NcoI-HF and XhoI and inserting HsCen2 as well as codon optimized PfCen1, ΔN-PfCen1, or PfCen2 sequences PCR amplified using primer pairs P24+25 and P26+27, P28+27, and P30+31 respectively. pET-28_PfCen1-GFP was generated by integrating a PfCen1 and GFP sequence PCR amplified using primer pairs P26+29 and P32+33 respectively. We generated pZE13d-C, a version of the bacterial expression vector pZE13d 45 with a C-terminal 6-His tag. For this purpose, pZE13d was digested with BamHI-HF and PstI-HF. A de novo sequence was formed by annealing oligo 3 and 4 over a 30 min temperature gradient from 95°C to 25°C and integrated via T4 ligase (New England Biolabs). pZE13d-C_PfCen3 and _PfCen4 were generated by digesting pZE13d-C with BamHI-HF and SalI-HF and integrating codon optimized PfCen3 and PfCen4 encoding sequences PCR amplified via primer pairs P34+35 and P36+37. All pZE13d derived bacterial vectors contained an ampicillin resistance cassette and a kanamycin resistance cassette for pET-28 derived vectors.

### *P. falciparum* blood culturing

All *P. falciparum* cell lines were maintained in human O+ erythrocyte cultures with a 2.5% haematocrit in RPMI 1640 medium supplemented with 0.5% AlbuMAX II Lipid Rich bovine serum albumin (Thermo Fisher Scientific), 25 mM Hepes (Sigma-Aldrich), 0.2 mM hypoxanthine (c.c.pro GMbH), and 12.5 µg/ml Gentamycin sulfate (Carl Roth) at pH 7.3 (“complete RPMI”). Cultures were incubated at 37°C in 3% CO2, 5% O2, and 90% humidity at 37°C. Parasitemia was maintained below 5% and determined via Hemacolor staining of blood smears (Sigma-Aldrich, Merck) using a Nikon Eclipse E100 microscope (Nikon Corporation) with 100x oil immersion objective. Depending on the application asexual cycle stages were synchronized to the ring stage by incubation of infected red blood cells (iRBCs) in a 25x volume of prewarmed 5% sorbitol (Sigma-Aldrich) solution at 37°C for 10-15 min, followed by a wash step in cRPMI (complete RPMI medium) before return to culture.

### P. falciparum transfection

For generation of transgenic parasite lines *P. falciparum* NF54 served as the acceptor strain for pARL plasmids and 3D7-DiCre (kindly provided by Anthony Holder and Mortiz Treeck) for pFIO plasmids 46,47. Plasmids were purified from 200 ml transformed *E. coli* overnight culture (37°C, 130 rpm) via the NucleoBond Xtra Midi Kit (Macherey-Nagel) and precipitated by supplementing 0.1x volume 3 M sodium acetate (Sigma-Aldrich, Merck, pH 5.3) and a 2x volume −20°C ethanol (VWR Chemicals). After overnight incubation at −20°C the DNA was pelleted at 17000g for 30 min at 4°C, washed with 70% ethanol, and air-dried under sterile conditions. For transfection 150 µl of iRBCs with a ≥ 4% ring stage parasitaemia were washed with prewarmed CytoMix [120 mM KCl, 0.15 M CaCl2, 2 mM EGTA, 5 mM MgCl2, 10 mM K2HPO4/KH2PO4, 25 mM Hepes, pH 7.6], then mixed with 100 µg purified plasmids in 30 µl TE buffer [10 mM Tris, 1 mM EDTA, pH 8.0] and 350 µl CytoMix. The mixture was transferred into a pre-cooled (4°C) Gene Pulser Electroporation cuvette (Bio-Rad) and electroporated using the Gene Pulser II system (Bio-Rad) at 310 kV, 950 µF, exponential mode. iRBCs were then transferred to 5 ml pre-warmed cRPMI for 30 min recovery at 37°C before centrifugation at 800g for 2 min and transfer to regular culture. Selection for episomal plasmids began 6 h post transfection and was henceforth maintained with 2.5 mM WR99210 (Jacobus Pharmaceutical Company).

### pFIO induction

Expression of the second pFIO ORF was induced in synchronous 3D7-DiCre_pFIO/pFIO+ parasite lines approximately 48 h prior to experiments during the schizont stage 48. 25 µl iRBCs were incubated in 200 µl cRPMI supplemented with 100 nM Rapamycin (Sigma-Aldrich, Merck) or 1% DMSO (Sigma-Aldrich, Merck) as a control for 4 h at 37°C. Afterwards cells were pelleted 1 min at 800 g for washing 4 times and cultured in 1 ml cRPMI without selection drug until further processing for imaging or FACS analysis.

### GFP quantification via flowcytometry

For flowcytometric quantification of GFP signal in induced 3D7-DiCre pFIO/pFIO+ strains 2.5 µl iRBCs were washed once with RT 0.9% NaCl solution (B. Braun GmbH) and stained with 5 µM SYTO 61 Red Fluorescent Nucleic Acid stain (Thermo Fisher Scientific) for 1 h at 37°C in 0.9% NaCl solution and washed once again afterwards at RT. Cells were analysed at RT using the FACSCanto II (BD Biosciences), first gating erythrocytes in the FSC-A/SSC scatter and then single cells in the FSC-H/FSC-A scatter. SYTO 61 signal was acquired using a 640 nm laser and 660/20 emission filter. Of the positive cells, i.e., iRBCs, multinucleated cells, i.e. schizonts, were selected and the share of GFP positive cells determined via a 488 nm laser and 530/30 nm emission filter, for which the gate cut-off was determined based on the uninduced DMSO control.

### Immunofluorescence assays

Immunofluorescence assay (IFA) of parasites for confocal and STED microscopy was performed as described previously in detail 49. In brief, parasites were seeded to concanavalin A (Sigma-Aldrich) coated µSlide 8-Well glass bottom dishes (Ibidi), fixed the next day with 4% PFA (Electron Microscopy Sciences) in PBS (Gibco, Thermo Fisher Scientific) for 20 min at 37°C, treated with 0.1% Triton X-100 (Sigma-Aldrich) and 0.1 mg/ml NaBH4 (Sigma-Aldrich) PBS solutions for permeabilization and quenching of free aldehyde groups. Blocking as well as incubation of primary and secondary antibodies (Table S3) was performed with 3% w/v Albumin Fraction V (Roth) in PBS. Hoechst 3342 (Thermo Fisher Scientific) was added during secondary antibody incubation at a dilution of 1:1000. Unbound antibodies were washed off with 0.5% Tween-20 (Roth) in PBS.

### ECCA frequency quantification

To quantify the frequency of ECCAs of PfCen1-GFP for different expression levels or mutants the respective seeded 3D7-DiCre_pFIO/pFIO+ strains were PFA fixed approximately 48 h post induction and an IFA for staining of DNA and tubulin was performed. In addition, GFP was stained using GFP-booster to preserve GFP signal with minimal off-target staining if the sample was analysed over multiple imaging sessions. Individual cells were acquired using the Leica Sp8 point laser scanning confocal microscope using HC PL APO CS2 63x/1.4 N.A. oil immersion objective. 128×128 pixel multichannel images were acquired sequentially using HyD detectors in the standard mode with a pixel size of 72.6 nm and z-interval of 0.3 µm with a pinhole of 1 airy unit. For 3D7-DiCre_pFIO-Cen1-GFP, _pFIO+_Cen1-GFP, and_pFIO+_ΔN-PfCen1-GFP 54, 60, and 51 early schizonts (cells with mitotic spindle and 1-5 nuclei) positive for GFP-foci were acquired in separate sessions. Foci outside the mitotic spindle poles were considered ECCAs. Analysis was performed using Fiji software 50. For induced 3D7-DiCre_pFIO-EFh-dead-Cen1-GFP no quantification was performed as no cells with foci were ever found. GFP signal density was quantified using Fiji by applying a threshold (1, 255) to the GFP signal to isolate the cellular signal from the background. Area and raw integrated density of each Z-slice was determined by via the multimeasure tool, added up, and divided.

### Proteostat staining of ECCA positive cells

To determine if ECCAs are amyloid aggregates live seeded 3D7-DiCre_pFIO+_Cen1-GFP were stained with 1:1000 Hoechst 3342 and 1:3000 Proteostat (Enzo Life Sciences) in cRPMI for 30 min at 37°C. Life cells were imaged at 37°C using the Leica SP8 microscope with the Lighting module, enabling automated adaptive deconvolution. 264×264 pixel images were acquired L HC PL APO CS2 63x/1.4 N.A. oil immersion objective was employed with GaAsP detectors, a pinhole of 0.6 airy units, a pixel size of 35.1 nm and z-stack of 7.28 µm at 130 nm intervals. PfCen1-GFP and Proteostat were excited with a 488 nm laser with GFP emission being measured between 490 to 550 nm and Proteostat emission at 600 to 700 nm. GFP bleed-through into the Proteostat range was negligible.

### STED microscopy

STED microscopy of NF54-pARL_Cen1-4-GFP was performed on an Expert Line STED system (Abberior Instruments GmbH) equipped with SLM based easy 3D module and an Olympus IX83 microscopy body, using an Olympus UPlanSApo 100x oil immersion objective/1.4 NA with a pixel size of 20 nm. STED images on fixed cells were acquired in RescueSTED mode (to avoid structural damage from the STED laser heating up the parasite’s hemozoin crystals) using the 590 and 640 nm excitation laser in line sequential mode with corresponding 615/20 and 685/70 emission filters. A 775 nm STED laser was employed at 10-15% intensity (maximum 3 mW) with a pixel dwell time of 10 µs. Regular confocal images were acquired in z-intervals of 300 nm, with 405 nm, 488 nm, 594 nm and 640 nm laser power being adjusted for each cell. Deconvolution of GFP staining signal was performed using Imspector Software (Abberior Instruments GmbH). For live-cell STED seeded NF54-pARL_Cen1-Halo were pre-treated with 1 µM SPY505-DNA (Spirochrome) and 1 µM MaP-SiR-Halo 36 for 3 h in cRPMI, which was replaced with cRPMI made from phenol red-free RPMI 1640 (PAN Biotech) prior to imaging. STED image series were acquired at 37°C in 1-30 second intervals on a single Z-slice using the same laser setup as above. Deconvolution was performed with Huygens professional software using express deconvolution with the standard template.

### Quantification of live PfCen1-GFP dynamics during schizogony

For long term live cell imaging of seeded 3D7-DiCre_pFIO/pFIO+-Cen1-GFP cells induced approximately 48 h prior were pre-incubated for 2 h at 37°C in an airtight 8-well dish with 0.5 µM SPY650-Tubulin in cRPMI made from phenol red-free RPMI 1640 that has been pre-equilibrated in 3% CO2, 5% O2 overnight. Imaging was performed at 37°C at the PerkinElmer UltraVIEW VoX microscope equipped with Yokogawa CSU-X1 spinning disk head and Nikon TiE microscope body with an Apo 60x/1.49 numerical aperture (NA) oil immersion objective and Hamamatsu C9100-23B electron-multiplying charge-coupled device (EM-CCD) camera with a pixel size of 59.8 nm. Images were acquired at multiple position using the automated stage and Perfect Focus System (PFS) in 5 min intervals. Multichannel images were acquired sequentially with 488 nm and 640 nm laser at 3% and 6% laser power as well as DIC. 8 µm stacks were acquired in 500 nm Z-intervals. For determination of the appearance of the first foci relative to the formation the first mitotic spindle the SPY650-tubulin signal was enhanced via deconvolution using the Fiji Deconvolutionlab2 with 10 iterations of the Richardson Lucy algorithm. PSF was generated using the PSF Generator plugin. Cells without visible focus at the start of imaging were visually analysed using Fiji. Mitotic spindle formation was identified by a strong increase in SPY650-tubulin foci signal and reduced mobility during the collapse of the hemispindle at the transition to a mitotic spindle.

For quantification of GFP signal within the cytoplasmic and foci fraction images were processed and analysed in a semi-automated workflow using ImageJ macros in the 64-bit 2.3.0 Fiji distribution of ImageJ 50. Image segmentation was carried out in 3D using version 1.3.2 of the Windows distribution of the image classification software ilastik 51. All macros are available on request. 3D timelapse GFP images were cropped to 80×80 pixel images containing individual iRBCs. Only iRBCs containing a single parasite were examined. To facilitate improved segmentation, image size was increased four times to 320×320 pixel using the adjust size function of ImageJ without interpolation. Segmentation of GFP foci was carried out on enlarged images using a pixel classification workflow in ilastik. With ilastik, a Random forest classifier utilizing all features up to and including a sigma of 3.5 in 3D was trained on two reference GFP timelapses via manual annotation. Subsequently, all 3D GFP timelapse images were batch processed using the same classifier. Segmentation files were exported as multipage tiff and reconstructed into 4D image sets matching the original 80×80 files by reordering stacks and decreasing the image size via the size adjust function of ImageJ without interpolation. Reconstructed segmentation images were then converted into binary masks containing only 0 (background) and 1 (object, here GFP foci) pixel values using the ImageJ math functions. Subsequently, signal intensities for GFP were measured in the following compartments over time: the foci, the whole parasite excluding the foci and the whole parasite including the foci. In parallel, the total area occupied by each compartment at a given time was also quantified. Results were collected as comma separated value (.csv) files and imported into Microsoft excel for analysis using an excel macro. To create masks for the whole parasite, images were segmented on background GFP values. Briefly, a median filter with a radius of 1 and a default threshold of 2200 (2200, 65535) were applied. To create binary parasite masks containing only 0 (background) and 1 (parasite) pixel values the thresholded image was divided by 255 using the ImageJ math functions. To create a mask of the whole parasite excluding the foci, the binary foci mask was inverted using the ImageJ math functions and the inverted foci mask multiplied with the parasite mask using the ImageJ image calculator function. The resulting binary mask contains only 0 (background and foci) and 1 (remaining parasite) pixel values. Areas were measured from binary segmentation masks by applying the multimeasure function on thresholded (1, 255) masks. To measure GFP signal intensities in different compartments, first background subtraction of 2000 was performed on the original 80×80 GFP images. The background subtracted image was then multiplied with one of the binary masks via the ImageJ calculator function, effectively removing any signal outside the compartment of interest. From the 4D image containing only the signal in the areas of interest, a summed z-projection was then created, from which signal intensity was measured as the raw integrated density using the ImageJ multimeasure function. Where indicated data of individual cells was normalized to the to the average value at the timepoints ranging from 10 min before and after mitotic spindle formation of that cell.

### Protein expression and purification

For the expression of PfCen1-6His variants as well as PfCen2-6His, and HsCen2-6His, chemically competent BL21-CodonPlus (De3) cells (Agilent Technologies) were transformed with the respective pET-28 vector via 42°C heat shock and plated overnight. Due to low transformation efficiency a single colony was picked for inoculation of an overnight pre-culture with 50 mg/L kanamycin (Roth), which was diluted 1:50 the next day in pre-warmed LB with kanamycin. For the expression of PfCen3-6His and PfCen4-6His chemically competent W3110-Z1 *E. coli* (Lutz and Bujard, 1996) were transformed with the respective pZE13d-C vector via 42°C heat shock and plated. The bacteria were scraped off and used to inoculate an expression culture at OD600 of 0.06 in pre-warmed LB with 50 mg/L ampicillin (Roth). These expression cultures were grown at 37°C, shaking at 130 rpm, until induction with 1 mM final IPTG (Thermo Scientific) once an OD¬600 of 0.5 for W3110-Z1 or 0.7 for BL21 cells was reached, after which incubation continued for 3 and 5 h respectively. All further steps were performed at 4°C. Bacteria were centrifuged and lysed via sonication in lysis buffer [3 mM β-Mercaptoethanol, 20 µg/ml DNaseI, 10 µg/ml Lysozyme, C0mplete protease inhibitor (Sigma-Aldrich), in PBS, pH 7.4]. Lysate was cleared via centrifugation at 17000 g for 20 min, supplemented with imidazole (Roth) to 10 mM, and the soluble fraction transferred to Ni-NTA beads (QIAGEN) in a gravity flow column. Beads were washed with a 10x volume of wash buffer [50 mM NaH2PO4H2O, 300 mM NaCl, pH 8.0] at 20 mM imidazole and protein eluted at 250 mM imidazole. Buffer was exchanged to 50 mM BisTris pH 7.1 via consecutive 2 h and overnight dialysis in a 200x volume using a 6-8 kDA cut-off tubing (Spectra/Por, 132650). Aliquots were snap-frozen in liquid nitrogen for storage at 80°C and slowly thawed on ice prior to experiments. Purity was determined via Coomassie gel. Concentrations were determined via Pierce 660 nm protein assay (Thermo Fisher) using pure bovine serum albumin (Thermo Fisher) as a standard. Purity was confirmed by loading 1.125 µg of protein on a 12-well 12% Mini-Protean TGX gel (Bio-Rad), performing SDS-PAGE electrophoresis and fixing the protein with 40% methanol, 10% acetic acid solution and staining with 0.25% Coomassie-blue (Merck) in 50% methanol, 10% acetic acid. Gels were documented with the Li-COR Odyssey CLx (Li-COR Biosciences) at 700 nm. Correct protein sizes were confirmed via LC-MS by the CFMP at the ZMBH Heidelberg, using a maXis UHR-TOF (Bruker Daltonics) coupled with nanoUPLC Acquity (Waters) tandem mass spectrometer for intact protein detection.

### LLPS quantification and imaging

To quantify LLPS over time 350 µl of protein sample solutions in 50 mM BisTris at pH 7.1 buffer were prepared in a 1.5 ml polystyrene cuvette (Sarstedt) on ice and transferred into a NP80 nanophotometer (Implen), which was blanked with buffer and preheated to 37°C. In case of PfCen1-GFP the protein solution itself was used for blanking due to absorption of GFP. The OD¬200-900 10 mm pathlength was then measured in 10 s intervals for 25 min, or 1 min intervals for up to 3 h in case of long-term measurements. At 450 s 3.5 µl 200 mM CaCl2 or MgCl2 solution was added to 2 mM (first 330 s not shown), and at 1170 s 7 µl 500 mM EDTA solution to 10 mM. To ensure even distribution of the solutions during addition into the cuvette they were pre-loaded into a 200 µl pipette tip, which was used to pipette sample solution up and down three times in between two measurements. To easily detect droplet fusion events, we used the following centrin concentration for the movies (PfCen1-6his, 200 µM. PfCen2-6his, 120 µM. PfCen3-6his, 193 µM. PfCen4-6his, 120 µM. HsCen2-6his, 200 µM). Otherwise, 30 µM was used or as indicated. 20 µl protein sample were imaged in non-binding PS µCLEAR 384 well plates (Greiner). For fluorescent imaging the 30 µM protein solutions were supplemented with 300 nM (100:1 molar ratio) NTA-Atto 550 (Sigma Aldrich, Merck). Movies were acquired at 37°C with a PerkinElmer UltraVIEW VoX microscope equipped with Yokogawa CSU-X1 spinning disk head and Nikon TiE microscope body with an Apo 60x/1.49 numerical aperture (NA) oil immersion objective and Hamamatsu sCMOS OrCA Flash 4.0 camera with a pixel size of 89.7 nm. Image series were taken in 1 s intervals using differential interference contrast (DIC) and 561 nm laser. For addition of 0.2 µl 200 mM CaCl2 solution or 0.4 µl 500 mM EDTA solution to the well the plate was briefly removed from the microscope for access. To assess co-condensation 150 µl of a 30 µM PfCen3-6His was imaged inside an untreated µSlide 8-Well glass bottom dishes (Ibidi), induced with 1.5 µl 200 mM CaCl2 solution, and supplemented during imaging with 20 µl PfCen1-GFP-6His directly into the well to a final concentration of 30 µM. GFP was excited using a 488 nm laser.

### Analysis of N-terminal IDRs in centrins

Likelihood of IDRs was estimated using the IUPred3 tool 52 (www.iupred3.elte.hu). Centrins with a ≥50% likelihood for N-terminal IDRs were considered positive. The phylogenetic tree was generated using ClustalOmega (www.ebi.ac.uk/Tools/msa/clustalo/), using default settings.

### Statistics

Statistical significance of the difference in the timepoint of PfCen1-GFP foci appearance between pFIO and pFIO+ (Figure 4E) was assessed via a two-tailed t-test with n=46 and 45 respectively.

## Acknowledgments

We thank: The Infectious Diseases Imaging Platform for imaging support (idip-heidelberg.org). PlasmoDB.org for their Plasmodium Informatics Resources. Core Facility for Mass Spectrometry & Proteomics (ZMBH). Nicolas Lardon and Kai Johnsson (Max Planck Institute for Medical Research) for providing the MaP-SiR-Halo dye. Anthony Holder and Moritz Treeck for the 3D7 DiCre strain. Marina Iocca for help with molecular cloning and protein expression. Friedrich Frischknecht, Sebastian Baumgarten, Daniel Gerlich, and Gautam Dey for critical discussion of the manuscript.

## Funding

We thank the German Research Foundation (DFG) 349355339, the Human Frontiers Science Program (HFSP) CDA00013/2018-C, and the Chica and Heinz Schaller Foundation for funding J.G. The Studienstiftung des Deutschen Volkes for funding Y.V. The German Research Foundation (DFG) 240245660 - SFB 1129 for funding M.G.

## Data availability statement

All data is contained within the manuscript and supplementary information. Additional information including ImageJ macros are available upon request.

## Conflict of interest disclosure

The authors declare that they have no competing interests.

## Notes

https://www.dropbox.com/sh/8c7bjy1uzixvmoh/AABIHMNK80mJXmNfrFPpiC3aa?dl=0

### Competing Interest Statement

The authors have declared no competing interest.

### Summary of Updates

Sections of the manuscript have been rewritten and reorganized. Data on co-condensation of centrins and supplemental movies have been added.

