## Supporting information for "Malaria parasite centrins assemble by Ca^2+^-inducible condensation"

Yannik Voß[1], Severina Klaus[1], Nicolas P. Lichti[1], Markus Ganter[1], Julien Guizetti[1]*
[1] Center for Infectious Diseases, Heidelberg University Hospital, Heidelberg, Germany.

**This PDF file includes:**

Figures S1 to S9

Tables S1 to S3

Captions for Movies S1 to S8

Movies can be downloaded under:

https://www.dropbox.com/sh/8c7bjy1uzixvmoh/AABIHMNK80mJXmNfrFPpiC3aa?dl=0

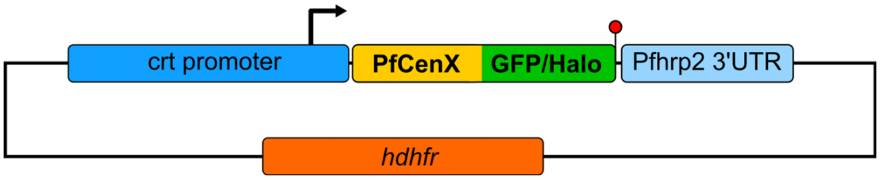


**Figure S1. Schematic map of pARL vector used for ectopic expression of tagged centrins.** The pARL vector contains a hDHFR cassette, conferring resistance to antifolates, and drives expression of a gene or fusion gene of choice from a weakened promoter of the *P. falciparum* chloroquine resistance transporter (Pfcrt).

A

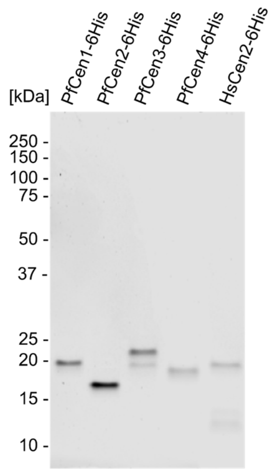


B

| | Expected size [kDa] | Major peak size [kDa] | Minor peaks size [kDa] |
|---|---|---|---|
| PfCen1-6His | 20.422 | 20.291<br>Met1 absent | |
| PfCen2-6His | 20.134 | 20.003<br>Met1 absent | |
| PfCen3-6His | 21.736 | 21.736 | 21.605<br>Met1 absent<br>20.085<br>additional band |
| PfCen4-6His | 20.926 | 20.926 | |
| HsCen2-6His | 20.561 | 20.430<br>Met1 absent | 11.255<br>12.545<br>additional bands |

C

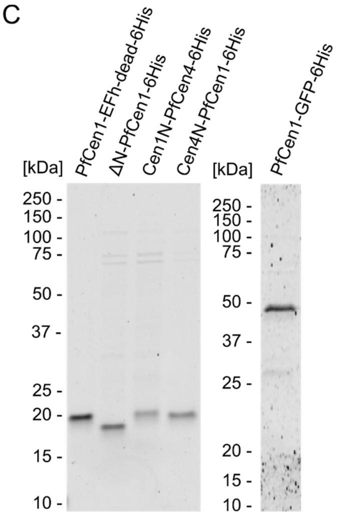


**Figure S2. Recombinantly produced centrins have full length size.** (A) Coomassie gels of wild type centrins recombinantly produced in bacteria and purified via their C-terminal His-tag shows slightly lower migration than expected for PfCen2-6His and 4 (B) Mass spectrometry analysis of native protein, however, confirms full length expression for all wild type centrins except for cleavage of Methionine 1 and occasional detection of degradation bands. (C) Coomassie gels of mutant versions of PfCen1 confirms proper expression and lower migration for deltaN-PfCen1 mutant.

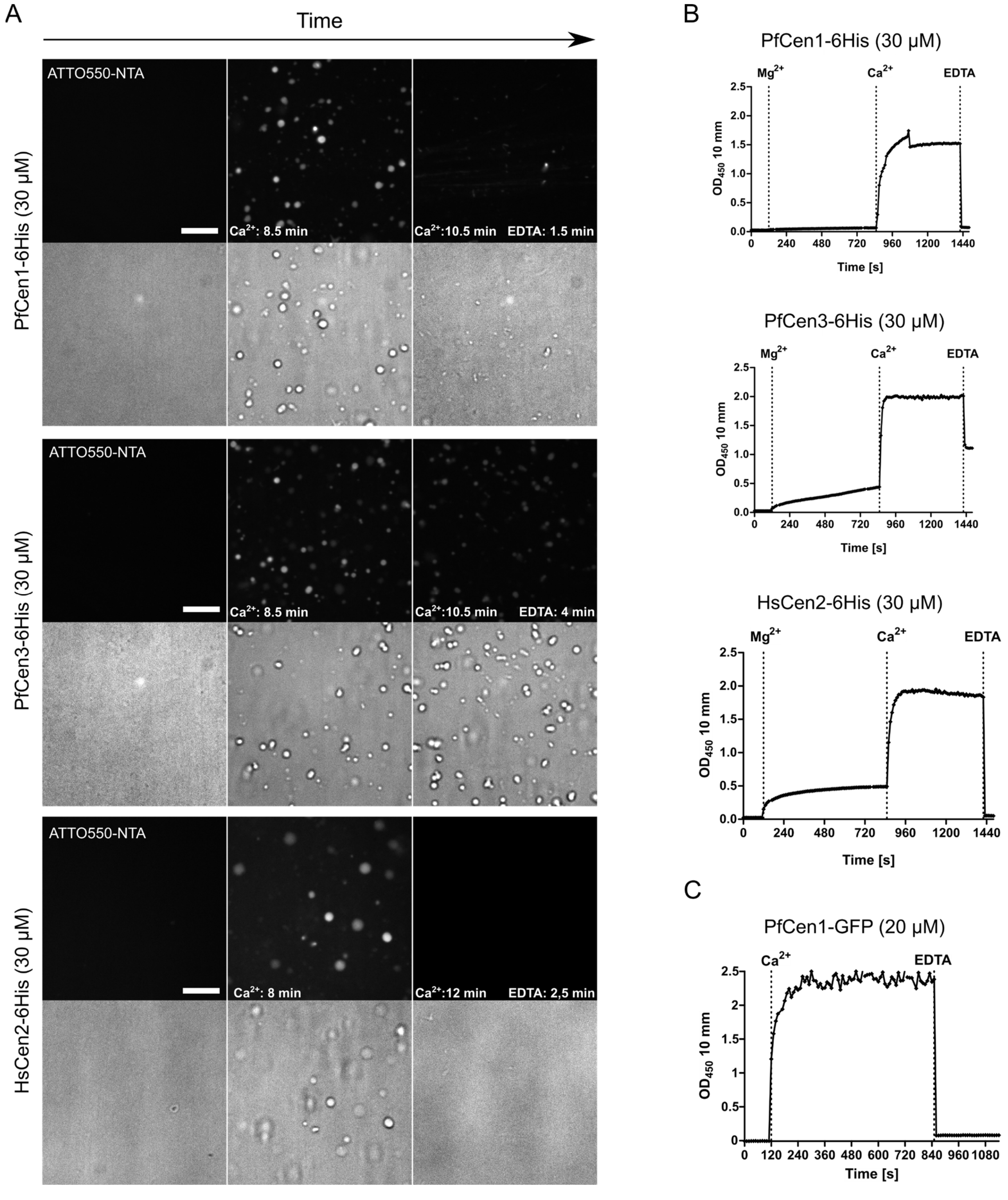


**Figure S3. Centrin phase-separation also occurs at lower concentrations and is calcium specific.** (A) Transmission light images of recombinant PfCen1, PfCen3, and HsCen2 centrin protein solutions at 30 µM concentration before, after calcium, and EDTA addition. Proteins were fluorescently labeled via their 6xHis tag using 300 nM NTA-Atto 550. Time stamps indicate time elapsed between calcium or EDTA addition and image acquisition. Observed droplets are more sparse than in highly concentrated solution but are still dynamic. (B) Turbidity assay using addition of magnesium followed by calcium addition at the same concentration indicates $Ca^{2+}$ specifically. and not bivalent cations per se, induces centrin LLPS. Conditions: 50 mM BisTris (pH 7.1), addition of $CaCl_2$ or $MgCl_2$ to 2 mM and EDTA to 10 mM, 37°C. (C) Assay as in (B) with recombinant PfCen1-GFP at 20 μM.

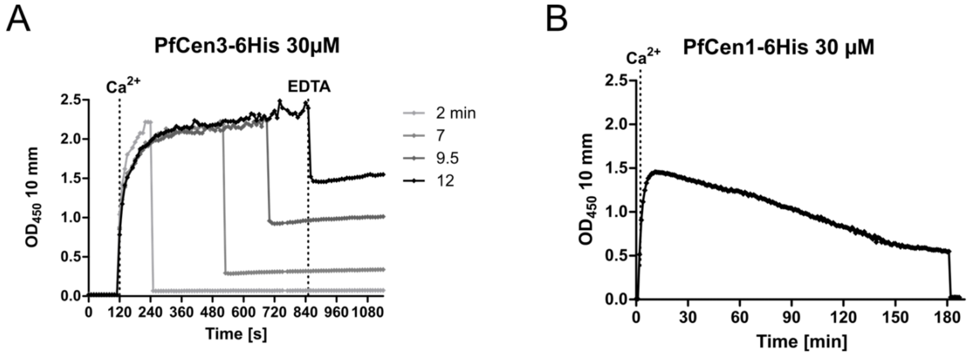


**Figure S4. Non-reversible condensate fraction of PfCen3 increases over time.** (A) Turbidity of PfCen3 solution with varying time points of EDTA addition after Calcium addition. (B) Turbidity of PfCen1 protein solution with highly delayed EDTA addition shows no irreversible fraction. Overall reduction of turbidity could be explained by protein droplets settling down. Conditions: 50 mM BisTris (pH 7.1), addition of $CaCl_2$ to 2 mM and EDTA to 10 mM, 37°C.

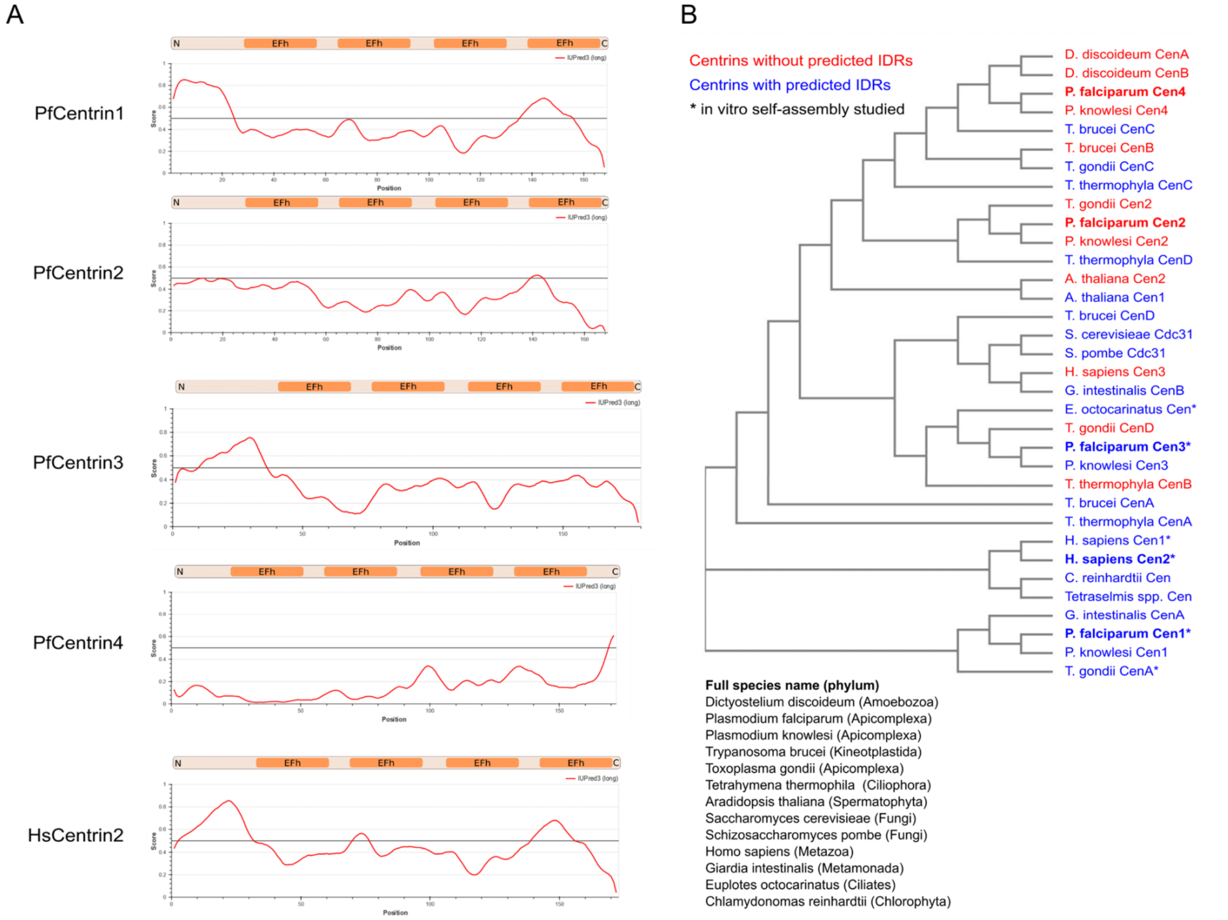


**Figure S5. Phase-separating centrins contain intrinsic disordered regions in their N-terminus.** (A) Using the IUpred 3.0 prediction software we identified a higher intrinsic disorder region probability (red line) in centrins undergoing LLPS in vitro. The highest probabilities are found in the N-terminus before the EFh domains. (B) Using Clustal Omega provided by the EMBL European Bioinformatics Institute we created a phylogenetic tree of centrins from multiple highly divergent eukaryotes, for several of which centrin self-assembly has been shown in vitro in this and previous studies (*). Using IUpred 3.0 we determined centrins with an IDR probability above the 0.5 threshold (blue) or without increased IDR probability (red). Centrins analyzed in this study are in bold.

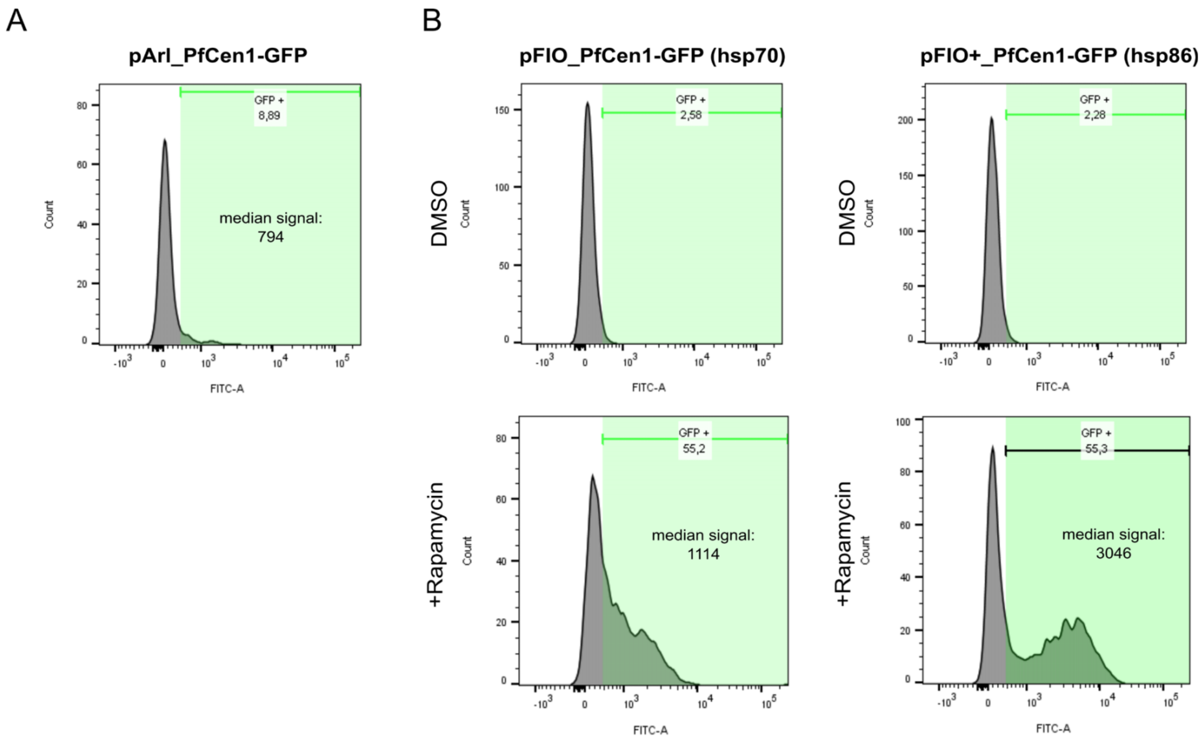


**Figure S6. pFIO constructs enable robust inducible overexpression of PfCen1-GFP in malaria parasites.** (A) Flow cytometry analysis of multinucleated malaria parasites stained with SYTO 61 as a DNA marker expressing PfCen1-GFP using the classical pARL construct only detects a small fraction of GFP-positive cells with a low median fluorescence intensity. (B) Fraction of GFP-positive parasites strongly increases upon induction of pFIO-Cen1-GFP transfected cells by Rapamycin addition. Mean GFP fluorescence intensity is much higher than in pARL whereas the hsp86 promoter generates even higher values than the hsp70 promoter fragment.

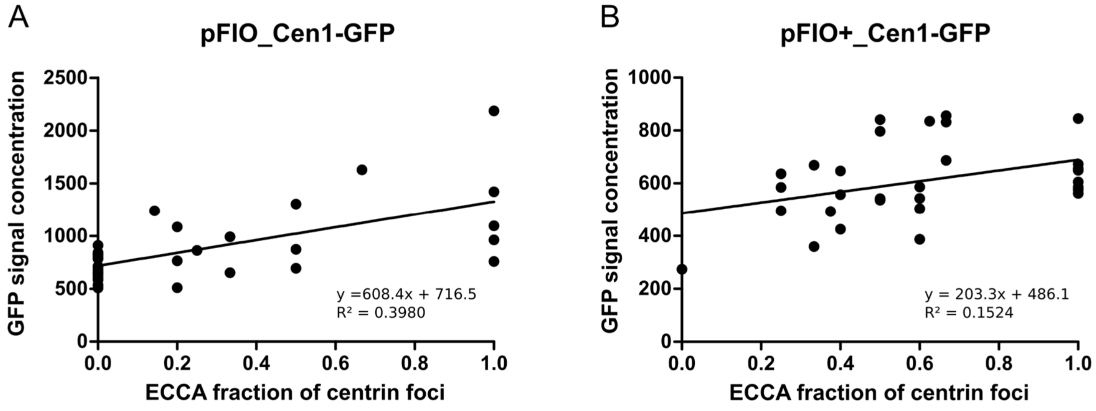


**Figure S7. ECCA appearance correlates with Cen1-GFP expression levels.** (A) Graph shows total arbitrary GFP fluorescence intensity of cells transfected with pFIO/pFIO+_Cen1-GFP one asexual cycle after induction plotted against the fraction of centrin foci, which are ECCAs per centriolar plaque associated centrin foci. Despite the low r-squared value statistical analysis indicate that the slope is significantly not zero ($p<0.0001$) and therefore positively correlated. (B) as in A but measured for pFIO+_Cen1-GFP expressing cells. Slope of linear regression is also significantly not zero ($p<0.033$). Due to the different expression levels different excitation laser settings had to be used for both parasite lines.

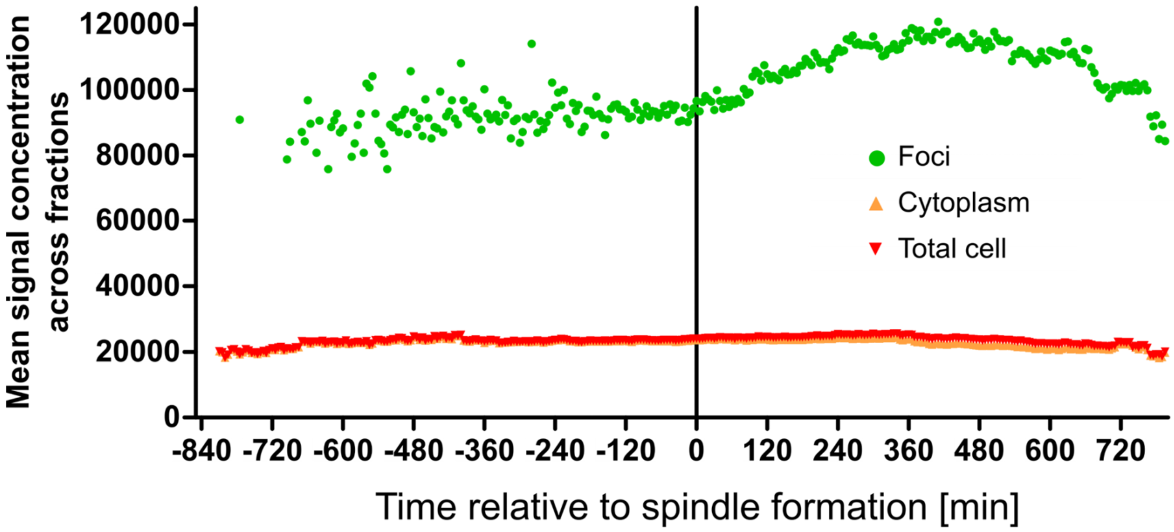


**Figure S8. Centrin signal concentration in foci is constant prior to spindle formation.** Mean PfCen1-GFP fluorescence intensity by segmented cellular region in induced parasites carrying pFIO+_PfCen1-GFP relative to mitotic spindle formation in time lapse movies. N $\leq$ 33 (see Supplementary Table S4 for details).

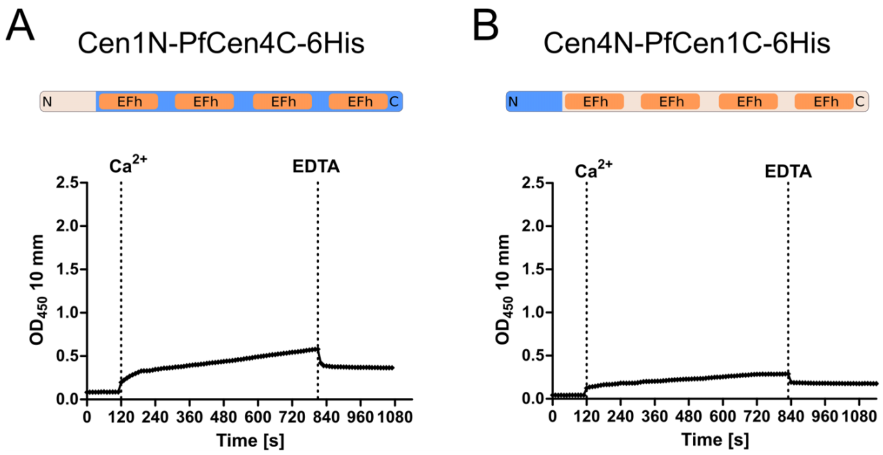


**Figure S9. PfCen1 N-terminus is essential but not sufficient for centrin phase separation.** (A) Turbidity of chimeric centrin fused from disordered N-terminus of PfCen1 and EFh domain containing part of PfCen4 during calcium and EDTA addition to protein solution. (B) Turbidity of chimeric centrin fused from N-terminus of PfCen4 and EFh domain containing part of PfCen1 during calcium and EDTA addition to protein solution. Conditions: 50 mM BisTris (pH 7.1), addition of $CaCl_2$ to 2 mM and EDTA to 10 mM, 37°C.

| # | Amplicon | Template | Dir. | Sequence |
|---|---|---|---|---|
| P1 | PfCen1 | NF54 cDNA | F | cgacccgggatggtaccATGAGCAGAAAAAATCAAACTATG |
| P2 | PfCen1 | NF54 cDNA | R | ttcttctcctttactcctaggAAATAAGTTGGTCTTTTTCATAATTC |
| P3 | PfCen2 | NF54 cDNA | F | cgacccgggatggtaccATGACCGATAACACAGCTG |
| P4 | PfCen2 | NF54 cDNA | R | ttcttctcctttactcctaggTAAGAAGCTTTTTTTGGTCATTATTG |
| P5 | PfCen4 | NF54 cDNA | F | cgacccgggatggtaccATGAACACAATGTTAATTAAGG |
| P6 | PfCen4 | NF54 cDNA | R | ttcttctcctttactcctaggTGAATCTGAATCAACATCAC |
| P7 | Halo+Linker | p238-actin-chromobody-halo* | F | gaaaaagaccaacttatttcctaggCCAACCACTGAGGATCTGT |
| P8 | Halo+Linker | p238-actin-chromobody-halo* | R | ccctcgaggaattcctgcaggtctggacatTTAACCGGAAATCTCCAGAGTAGAC |
| P9 | PfHsp70 5' UTR | NF54 gDNA | F | cactatagaatactcgcggccgcCGCATAAATATCTGGTGAAATACAAAC |
| P10 | PfHsp70 5' UTR | NF54 gDNA | R | atgtatgctatacgaagttatgaaCCTTTTGCACTAGCCAATTTTTC |
| P11 | hDHFR+GGGS linker | pARL-Cen3-GFP | F | cttcgtatagcatacattatacgaagttatATGCATGGTTCGCTAAAC |
| P12 | hDHFR+GGGS linker | pARL-Cen3-GFP | R | aatctattattaaataaatggatccacctccaccATCATTCTTCTCATATACTTC |
| P13 | hDHFR+GGGS linker | pARL-Cen3-GFP | R | TCTTCTCCCTTAGATACGG |
| P14 | mCherry | p3xNLS-mCherry-hsp86-BSD | F | gaatgatggtggaggtggatccGTATCTAAGGGAGAAGAG |
| P15 | mCherry | p3xNLS-mCherry-hsp86-BSD | R | gtataatgtatgctatacgaagttatgtcgacttaTGTAGAGTGTCTACCTTC |
| P16 | GFP | pARL-Cen3-GFP | F | gtatagcatacattatacgaagttatacgcgtATGGTTAGTAAAGGAGAAGAACTTTTC |
| P17 | GFP | pARL-Cen3-GFP | R | ctattattaaataaatgctcgagTTATTTGTATAGTTCATCCATG |
| P18 | PbDHFR 3'UTR | PbANKA gDNA | F | gtagacactctacataagtcgacGATATGGCAGCTTAATGTTC |
| P19 | PbDHFR 3'UTR | PbANKA gDNA | R | atgctatacgaagttat G ATATCGAAATTGAAGG |
| P20 | PfHsp86 5'UTR | NF54 gDNA | F | cactatagaatactcgcggccgcGGGAATTTCCTTATAAGATCTTTC |

| | | | | |
|---|---|---|---|---|
| P21 | PfHsp86 5'UTR | NF54 gDNA | R | atgtatgctatacgaagttATTTTATTCGAAATGTGGGAAG |
| P22 | Cen1 | pARL-Cen1-GFP | F | acattatacgaagttatacgcgtATGAGCAGAAAAAATCAAAC |
| P23 | ΔN-PfCen1 | pARL-Cen1-GFP | F | acattatacgaagttatacgcgtATGGAATTAAATGAAGAACAAAAATTAG |
| P24 | HsCen2 | human cDNA | F | actttaagaaggagatataccATGGCCTCCAACTTTAAGAAGG |
| P25 | HsCen2 | human cDNA | R | agtggtggtggtggtggtgcccATAGAGGCTGGTCTTTTTCATG |
| P26 | PfCen1 (e. coli) | oligo 1 | F | actttaagaaggagatataccATGTCTCGTAAAAACCAAACC |
| P27 | PfCen1 (e. coli) | oligo 1 | R | agtggtggtggtggtggtgcccGAATAAGTTAGTTTTTTTCATAATGCG |
| P28 | ΔN-PfCen1 (e. coli) | oligo 1 | F | tttaagaaggagatataccatgGAGCTGAATGAGGAACAAAAG |
| P29 | PfCen1 (e. coli) | oligo 1 | R | ttctcctttactcctaggGAATAAGTTAGTTTTTTTCATAATGCG |
| P30 | PfCen2 (e. coli) | oligo 2 | F | actttaagaaggagatataccATGACAGATAACACTGCC |
| P31 | PfCen2 (e. coli) | oligo 2 | R | agtggtggtggtggtggtgcccCAGGAAACTTTTCTTGGTC |
| P32 | GFP | pARL-Cen3-GFP | F | CCTAGGAGTAAAGGAGAAGAAC |
| P33 | GFP | pARL-Cen3-GFP | R | agtggtggtggtggtggtgcccTTTGTATAGTTCATCCATGCC |
| P34 | PfCen3 (e. coli) | oligo 3 | F | ttaaagaggagaaaggatccATGATTAACCGTAAGAGC |
| P35 | PfCen3 (e. coli) | oligo 3 | R | gtggtgatgatggtgATATAAGGATGTCTGCTTC |
| P36 | PfCen4 (e. coli) | oligo 4 | F | ttaaagaggagaaaggatccATGAATACTATGCTGATCAAAG |
| P37 | PfCen4 (e. coli) | oligo 4 | R | gtggtgatgatggtgAGAGTCGCTGTCAACGTC |

**Table S1. Primers used in this study.** All primers used for amplification of the various construct fragments used for molecular cloning as described above are listed here. Homology regions for Gibson Assembly are in lowercase, PCR binding sequences are capitalized. * Kindly provided by Dr. Friedrich Frischknecht.

| Oligos | Encodes | Sequence |
|---|---|---|
| oligo 1 | EFh-dead PfCen1 aspartate 37, 73, 110, 146 mutated to alanine | acattatacgaagttatacgcgtATGAGCAGAAAAAATCAAACTATGATAAGGAACCCTAATCCTCGAAGTAAAAGAAACGAATTAAATGAAGAACAAAAATTAGAAATTAAGGAAGCTTTTGATTTATTTGCTACAAACGGCACTGGAAGAATTGATGCAAAAGAATTAAAAGTTGCAATGAGAGCTTTAGGATTCGAACCAAAAAAAGAAGATATAAGGAAAATAATATCTGATGTTGCTAAAGATGGATCTGGTACAATTGACTTTAATGACTTTTTAGACATTATGACAATAAAAATGAGTGAAAGAGATCCTAAGGAGGAAATACTAAAAGCTTTCCGATTATTTGCTGATGATGAGACTGGAAAAATCTCCTTCAAAAATTTAAAACGTGTAGCAAAAGAACTTGGAGAGAATATTACTGATGAGGAAATTCAAGAAATGATAGACGAAGCAGCCAGAGATGGAGACGGAGAAATTAATGAAGAAGAATTTATGAGAATTATGAAAAAGACCAACTTATTTcctaggagtaaaggagaagaa |
| oligo 2 | HsCen2 codon optimized for *P. falciparum* | acattatacgaagttatacgcgtATGGCAAGTAATTTTAAAAAAGCAAATATGGCAAGTTCAAGTCAAAGAAAAAGAATGAGTCCAAAACCAGAATTAACAGAAGAACAAAAACAAGAAATTAGAGAAgcaTTTGATTTAttcGATGCAGATGGAACTGGAACAATAGATGTAAAAGAATTAAAAGTAGCAATGAGAGCATTAGGATTTGAACCTAAAAAAGAAGAAATTAAAAAAATGATAAGTGAAATAGATAAAGAAGGAACTGGAAAAATGAATTTTGGAGATTTTcttACAGTAATGACACAAAAAATGAGTGAAAAAGATACAAAAGAAGAAATATTAAAAGCTTTTAAACTTTTTGATGATGATGAAACTGGAAAAATTtccTTTAAAAATTTAAAAAGAGTAGCTAAAGAActtGGAGAAAATTTAACAGATGAAGAATTACAAGAAATGATAGATGAAGCAGATAGAGATGGAGATGGAGAAGTATCAGAACAAGAATTTTTAAGAATTATGAAAAAAACATCATTATATcctaggagtaaaggagaagaa |
| oligo 3 | 6His | gatcCATACGCGTCGACACCATCATCACCACCACTAACtgca |
| oligo 4 | 6His | GTTAGTGGTGGTGATGATGGTGTCGACGCGTATG |
| oligo 5 | PfCen1 codon optimized for *E. coli* | catcaccaccacatcgatATGTCTCGTAAAAACCAAACCATGATTCGTAATCCGAATCCGCGTAGCAAACGCAACGAGCTGAATGAGGAACAAAAGCTTGAAATTAAAGAAGCGTTCGATCTTTTTGACACAAATGGTACAGGGCGTATCGACGCAAAAGAGCTTAAAGTGGCAATGCGCGCATTGGGGTTTGAACCAAAAAAGGAGGACATCCGTAAAATTATTTCAGATGTCGATAAAGACGGCTCTGGGACGATTGATTTCAATGACTTTTTAGACATTATGACCATTAAAATGTCGGAACGCGATCCCAAAGAGGAAATTCTTAAAGCGTTTCGTTTATTCGACGACGACGAAACCGGAAAAATTTCCTTCAAAAACCTGAAGCGCGTAGCAAAAGAGCTTGGAGAAAATATCACGGACGAGGAAATTCAAGAGATGATTGATGAAGCGGACCGCGATGGAGATGGAGAAATTAACGAGGAAGAGTTCATGCGCATTATGAAAAAAACTAACTTATTCTAActgcagccaagcttaatt |

| | | |
|---|---|---|
| oligo 6 | PfCen2<br><br>codon optimized for *E. coli* | catcaccaccacatcgatATGACAGATAACACTGCCGTGCGCCGCCGTTATGAAAAATCTTTCACGGAACGTCCGGGTTTACGGAGGATGAGATTGAGGAAATTCGTGAAGCCTTTAACCTTTGGACACCGACGGGACGGGGACCATCGATCCTAAGGAAATTAAATGTGCCATGCAATCGCTTGGCTTAGATGCAAAGAATCCCATGATTTTCCGCATGATCGCGGACCTGGAAAAAGACGGCTATTCCTCCATCGATTTTGAAGAATTCATGGAGGTGATTACAAGTAAGTTGGGCAACAAGGACACGCGTGAGGGCATTCAGCGCATCTTCAACTTATTTGATGACGATAAAACAGGTTCGATCTCTCTTAAAAACTTAAAACGTGTTGCTAAAGAGTTAGGCGAAACATTAACGGACGAAGAATTGCGTGATATGATTGATCGTGCCGATTCGAAGGGGGAGGGCGAAATCAGCTTCGAGGACTTTTACACTATTATGACCAAGAAAAGTTTCCTGTAActgcagccaagcttaatt |
| oligo 7 | PfCen3<br><br>codon optimized for *E. coli* | catcaccaccacatcgatATGATTAACCGTAAGAGCGAATCTGTCAGCTACACGCCTCGTGCAATCACTAACCGCCCTATTAGTAGCAATCGCCGTCGCGGGCGCAACGAAATTACAGACGAGCAGAAAAATGAAATTAAGGAAGCGTTTGATCTTTTCGATACAGAGAAAACCGGCAAAATCGACTATCATGAGCTGAAGGTTGCCATCCGTGCATTAGGATTCGATATTAAGAAAGCGGACGTTTTAGACTTGATGCGCGAATACGACAAGACCAACAGTGGGCATATCGATTATAACGATTTCTTGACATTATGACTCAAAAGATTTCTGAGCGTGACCCAACGGAAGAAATTATTAAGGCTTTTAAGCTTTTCGATGACGACGATACTGGGAAAATCTCCTTGAAGAATTTGCGTCGTGTTTCGCGTGAGCTGGGTGAAAATTTAAGCGATGACGAGTTACAAGCTATGATCGACGAATTCGACAAAGACATGGATGGGGAAATCTCTCAAGAGGAATTCTTGTCAATCATGAAGCAGACATCCTTATATTAActgcagccaagcttaatt |
| oligo 8 | PfCen4<br><br>codon optimized for *E. coli* | catcaccaccacatcgatATGAATACTATGCTGATCAAAGATAACATTAACATCTCAATTAACGAGGATGTAGAAAAAGAACTGTACGAATGCTTTTCACTGTTTGACACGAACAAGTGCGGCTACATTGATATTCGTGAGTTTTACTTTGCCTTGAAATCCTTGGGCCTGAACTTCAAGAAGGAGCAGGTTAAAAATATTTTTTTGGACATCAAGAAAGATATTGACGAGAAATTGAATTTTGACGAATTTTTTGATATCGCAACGAAATATCTGCATACCCGCTATAATGACGACGAAATGGACCAAATGTTCTCATTGTTTGACCCCAACGATACTGGTAAAATTACACTTCAGGCTCTGCGTAAGGTTTGCACTGACATCGGGGAGAATATCTCGGATACCGAGCTGAATAACATGATCCACTTCGCAGACAAAAACAATGATAAAGTCATCGATAAAAACGAGTTCAAAAAGGTGCTTCTTTGCTCCTGGAAGAATGATCCGTTATCGGACGTTGACAGCGACTCTTAActgcagccaagcttaatt |
| oligo 9 | EFh-dead PfCen1 aspartate 37, 73, 110, 146 | actttaagaaggagatataccATGTCTCGTAAAAACCAAACCATGATTCGTAATCCGAATCCGCGTAGCAAACGCAACGAGCTGAATGAGGAACAAAAGCTTGAAATTAAAGAAGCGTTCGATCTTTTTGCCACAAATGGTACAGGGCGTATCGACGCAAAAGAGCTTAAAGTGGCAATGCGCGCATTGGGGTTTGAACCAAAAAAGGAGGACATCCGTAAAATTATTTCAGATGTCGCGAAAGACGGCTCTGGGACGATTGATTTCAATGACTTTTTAGACATTATGACCATTAAAATGTCGGAACGCGATCCCAAAGAGGAAATTCTTAAAGCGTTTCGTTTATTCGCCGACGACG |

| | | |
|---|---|---|
| | mutated to alanine, E. coli codon | AAACCGGAAAAATTTCCTTCAAAAACCTGAAGCGCGTAGCAAAAGAGCTTGGAGAAAATATCACGGACGAGGAAATTCAAGAGATGATTGATGAAGCGGCCCGCGATGGAGATGGAGAAATTAACGAGGAAGAGTTCATGCGCATTATGAAAAAAACTAACTTATTCgggcaccaccaccaccaccact |

**Table S2. Oligos used in this study.** All listed oligos used for molecular cloning were ordered from Genestring (except 6His-tags).

| Antibody | Species | Dilution | Source |
|---|---|---|---|
| Anti-alpha-tubulin B-5-1-2, monoclonal | mouse | 1:500 | Sigma-Aldrich (T5168) |
| Anti-PfCentrin3, polyclonal | rabbit | 1:500 | Simon et al. 2021 |
| Anti-GFP, ABfinity Monoclonal | rabbit | 1:50 | Thermo Fisher (G10362) |
| Anti-mouse-Atto 594 | goat | 1:1000 | Sigma-Aldrich (76085-1ML-F) |
| Anti-mouse-Atto 647 | goat | 1:1000* | Sigma-Aldrich (50185-1ML-F) |
| Anti-rabbit-Atto 594 | goat | 1:1000* | Sigma-Aldrich (77671-1ML-F) |
| GFP-Booster_Atto 488 | - | 1:200 | Chromotek (gba488-100) |

**Table S3. Antibodies used in this study.** List of antibodies with information of species, final dilution and source with order number. Starred (*) are dilutions for confocal IFAs. For STED microscopy concentration was increased to 1:200.

**Movie S1. Some PfCen1-Halo foci display round fluid-like dynamics.** STED time lapse movie at 1 s time interval of centriolar plaque region of parasites expressing PfCen1-Halo labeled with MaP-SiR-Halo dye (green) DNA stained with SPY505-DNA. Scale bar, 100 nm. Linear bleaching correction was applied using Fiji.

**Movie S2. Some PfCen1-Halo foci display stretched fluid-like dynamics.** STED time lapse movie at 1 s time interval of centriolar plaque region of parasites expressing PfCen1-Halo labeled with MaP-SiR-Halo dye (green) DNA stained with SPY505-DNA. Scale bar, 100 nm. Linear bleaching correction was applied using Fiji.

**Movie S3 PfCen1 displays LLPS upon calcium addition in vitro.** Transmission time lapse imaging of recombinant PfCen1-6His protein solution at 200 µM concentration displays wetting and fusion events rapidly after calcium addition. 50 mM BisTris (pH 7.1), addition of $CaCl_2$ to 2 mM and EDTA to 10 mM, 37°C.

**Movie S4 PfCen3 displays LLPS upon calcium addition in vitro.** Transmission time lapse imaging of recombinant PfCen3-6His protein solution at 193 µM concentration displays wetting and fusion events rapidly after calcium addition. 50 mM BisTris (pH 7.1), addition of $CaCl_2$ to 2 mM and EDTA to 10 mM, 37°C.

**Movie S5 HsCen2 displays LLPS upon calcium addition in vitro.** Transmission time lapse imaging of recombinant HsCen2-6His protein solution at 200 µM concentration displays wetting and fusion events rapidly after calcium addition. 50 mM BisTris (pH 7.1), addition of $CaCl_2$ to 2 mM and EDTA to 10 mM, 37°C.

**Movie S6 PfCen3 droplets are not fully resolved by EDTA and display more solid-like properties.** Transmission time lapse imaging of recombinant PfCen3 protein solution at 193 µM concentration after calcium and EDTA addition does not dissolve droplets. Remaining spheres don't display fusion or wetting anymore. 50 mM BisTris (pH 7.1), addition of $CaCl_2$ to 2 mM and EDTA to 10 mM, 37°C.

**Movie S7. PfCen1-GFP overexpression causes ECCA formation.** Live cell spinning disk time lapse movie at 5 min time interval of parasite overexpressing PfCen1-GFP (green) from pFIO+ labeled with SPY650-Tubulin (magenta) during entry into schizogony at 37°C. Image dimensions are 15 x 15 µm. Left frame only shows microtubule signal. Maximum intensity projection.

**Movie S8. ECCAs frequently form at centriolar plaques before detaching.** Short live cell spinning disk time lapse movie at 5 min time interval of schizont parasite overexpressing PfCen1-GFP (green) from pFIO+ labeled with SPY650-Tubulin (magenta) showing detachment of the brightest foci from the spindle. Image dimensions are 15 x 15 µm. Left frame only shows microtubule signal. Maximum intensity projection.
